# Predicting diabetes second-line therapy initiation in the Australian population via timespan-guided neural attention network

**DOI:** 10.1101/529933

**Authors:** Samuele Fiorini, Farshid Hajati, Annalisa Barla, Federico Girosi

## Abstract

**Introduction:** The first line of treatment for people with diabetes is metformin. However, over the course of the disease metformin may fail to achieve appropriate glycemic control, and a second-line therapy becomes necessary. In this paper we introduce `Tangle`, a timespan-guided neural attention model that can accurately and timely predict the upcoming need for a second-line diabetes therapy from administrative data in the Australian adult population. The method could be used to design automatic therapy review recommendations for patients and their providers without the need to collect clinical measures.

**Data:** We analyzed seven years of deidentified records (2008-2014) of the 10% publicly available linked sample of Medicare Benefits Schedule (MBS) and Pharmaceutical Benefits Scheme (PBS) electronic databases of Australia.

**Methods:** By design, `Tangle` can inherit the representational power of pre-trained word embedding, such as GloVe, to encode sequences of claims with the related MBS codes. The proposed attention mechanism can also natively exploit the information hidden in the timespan between two successive claims (measured in number of days). We compared the proposed method against state-of-the-art sequence classification methods.

**Results:** `Tangle` outperforms state-of-the-art recurrent neural networks, including attention-based models. In particular, when the proposed timespan-guided attention strategy is coupled with pre-trained embedding methods, the model performance reaches an Area Under the ROC Curve of 90%, an improvement of almost 10 percentage points over an attentionless recurrent architecture.

**Implementation:** `Tangle` is implemented in Python using Keras and it is hosted on GitHub at https://github.com/samuelefiorini/tangle.

## Introduction

Diabetes affects around 1.2 million of Australians aged 2 years and over. In the last two decades, the prevalence of the disease almost doubled, reaching 5.1% of the population in 2015 ^1^. In the same year, 85% of the Australians with diabetes reported a Type 2 Diabetes Mellitus (T2DM) diagnosis. This type of disease is particularly worrisome as it is the leading cause of more than half of the diabetes-related deaths of 2015 [1]. In order to reach glycemic control in T2DM subjects, Diabetes Australia recommends dietary changes and physical exercise along with administration of metformin, if needed [2]. When metformin is not sufficient anymore to achieve good glycemic control, second-line medications should be added [3]. Failing to do so will lead to worsening conditions and therefore it is important to identify those patients who should be targeted for therapy change, so they can be monitored closely.

Thanks to recent advances in the field of machine learning it is becoming possible to design algorithms that exploit medical records to predict and identify those patients who benefit from specific interventions [4].

In this paper we describe a predictive algorithm that looks at the administrative medical records history of a patient and estimates the likelihood that they will need second-line medication in the next future. This method could be used to design an automatic system for patients and/or their providers that notifies them that a change in therapy might be worth considering. From a machine learning point of view this means that we build a classifier where the samples are sequences of medical events and the binary labels identify subjects that added a second-line medication.

The medical events we consider in this paper are any of the events reported for administrative purposes in the Medicare Benefits Schedule (MBS), that records the utilization of primary care services such as visits to GPs and specialists, diagnostic and pathology testing as well as therapeutics procedures. While using actual clinical records seems an appealing, albeit more complex, option and might results in better predictions, we have not considered it because an integrated system of health records has not been implemented yet at national level. MBS records, instead, are not only routinely collected at federal level for administrative purposes, but are also, to some extent, available for data analysis.

## Background

In this paper we focus on learning a classification function for sequences, *i.e.* ordered lists of of events, that are encoded by symbolic values [5]. A major challenge with this type of data is how to map them in a numerical representation suitable to train a classification model. Standard vector representations, adopted for instance in natural language processing, can be either *dense* (*i.e.* most of the elements are nonzero) or *sparse* (*i.e.* with only few nonzero elements). A popular sparse representation method for symbolic elements, or categorical features, is called One-Hot-Encoding (OHE) and consists in directly mapping each symbolic element to a unique binary vector [6]. Although frequently used, this representation acts at a local level and it is therefore necessary to adopt some feature aggregation policy to achieve a global representation of a given input sequence. Another sparse representation strategy is multidimensional Bag-of-words (BOW), where each dimension represents the number of occurrences of a given *n*-gram in the sequence [7].

Nowadays, *word embeddings* are the most popular dense representation for sequence learning problems. In this approach, to each element **w**_*i*_ of the sequence **s**(*i.e.* word of the document) one associates a real-valued dense vector 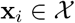. The semantic vector space 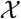 is designed to have “interesting” properties: *e.g.* neighboring vectors may correspond to words having similar meaning or sharing similar contexts. The two most popular word embeddings models proposed in literature are called word2vec [8] and Global Vectors for Word Representation (GloVe) [9].

Once a suitable encoding strategy is defined, a machine learning problem can be posed. In this context, standard sequence classification models can be linear, *e.g.* Logistic Regression (LR) and Support Vector Machines [10], or nonlinear, *e.g.* Random Forests [11] and Boosting [12]. These approaches usually are not as computationally expensive as other methods such as deep learning techniques and can be used in combination with feature selection schemes to promote interpretability of the results [13]. However, this class of techniques suffer from a major drawback: *i.e.* their predictive performance is *heavily* influenced by the discriminative power of the adopted sequence representation.

In the recent past, deep learning methods showed remarkable performance in solving complex prediction tasks, such as visual object and speech recognition, image captioning, drug-discovery and so on [14]. In the plethora of deep learning models, Recurrent Neural-Networks (RNN) [14] is the class of architectures specifically designed to work with sequential inputs. They consecutively process each element keeping a hidden state vector that can memorize information on the past history. Although designed to learn long-term dependencies, empirical evidence show that vanilla RNN fail in this task. On the other hand, Long Short-Term Memory (LSTM) networks [15], a particular class of RNN, are specifically designed to solve this issue. LSTMs have special memory cells that can work as information accumulator together with a system of input, output and forget gates. These networks empirically showed that they can deal well with both short and long-time relationship among the elements of input sequences. RNN, and deep learning models in general, can also easily inherit the representational power of pre-trained word embeddings, heavily increasing their classification performance [6]. A schematic representation of how RNN-based models can be used to solve a sequence classification task is presented in Fig. 1.

**Fig 1.**
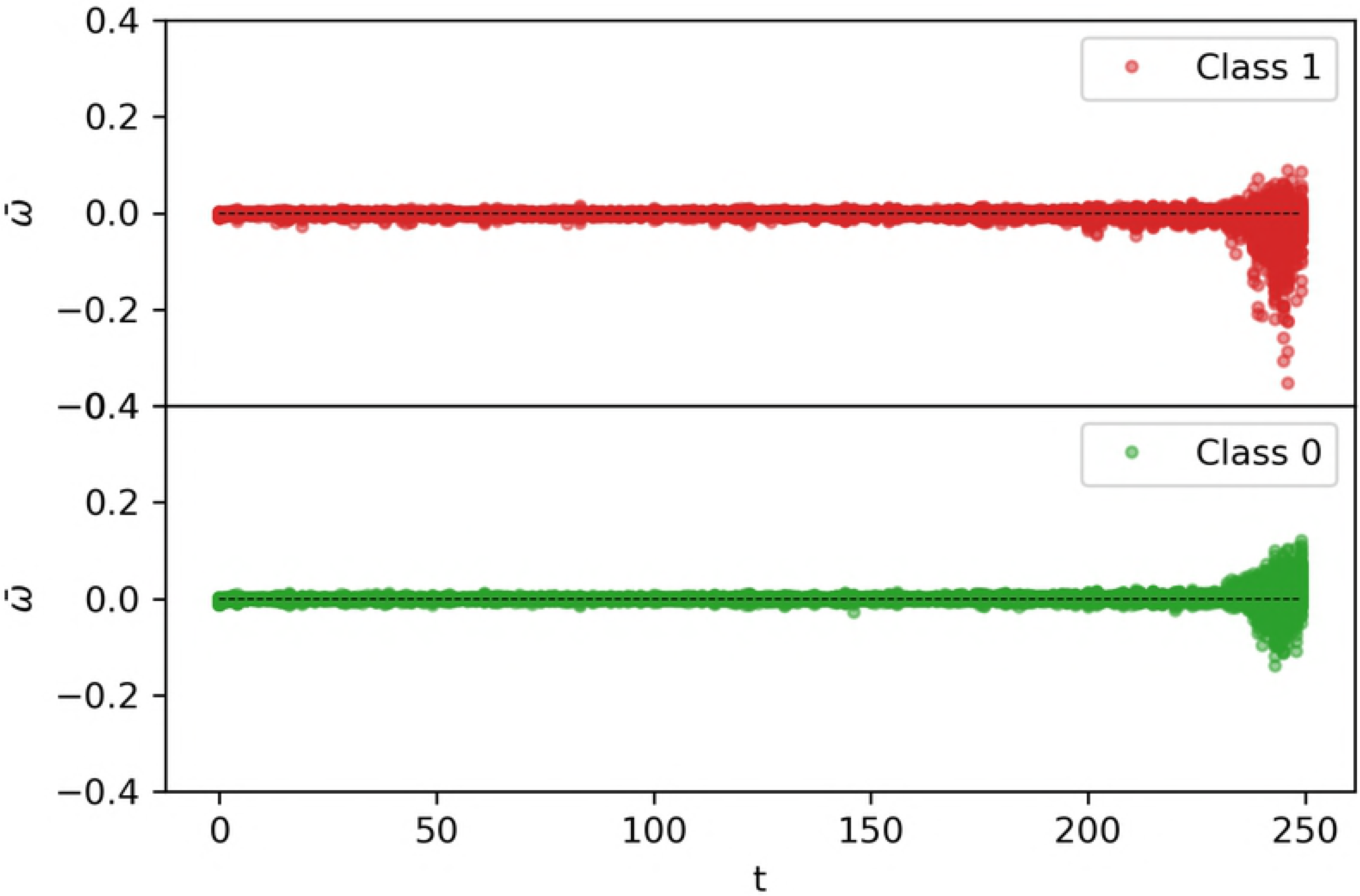
LSTM for sequence classification. A visual representation of a simple bidirectional LSTM for sequence classification model. This architecture is used in this work for the sake of comparison, and it is referred to as *baseline*. In this work we adopted LSTM recurrent cells, in order to exploit their ability to learn long-time relationship in the sequences. However, similar architectures can be devised with vanilla RNN, Gated Recurrent Units (GRU) [17] or other types of temporal architectures.

Two major shortcomings of these architectures is that: (i) in order to achieve their top performance they need to be trained on large datasets, hence requiring high computational time and (ii) when applied in health care-related settings the learned representations hardly align with prior (medical) knowledge [16]. For a comprehensive overview of the most widely adopted deep learning models see [14] and references therein.

Throughout this paper, real-valued variables are indicated with lowercase letters (*e.g. a*), unidimensional vectors with lowercase bold letters (*e.g.* **a**) and matrices, or tensors, with capital letters (*e.g. A*). To avoid clutter, sample subscripts are omitted where not strictly needed.

### Neural attention mechanism

Neural attention [18] is a recently proposed strategy to promote interpretability and to improve prediction performance of deep learning methods for document classification [19], machine translation [18], prediction from sequential Electronic Health Record (EHR) [16, 20, 21] and so on. The intuition behind attention mechanism is that not all elements of the sequence are equally relevant for the prediction task and that modeling their interactions helps to find the most relevant patterns.

Neural attention mechanism can be seen as a strategy to find *weights* (*α*) that can emphasize events occurring at some point in the sequence, with the final aim to improve the prediction performance. A possible adopted solution to find such weights is via Multi-Layer Perceptron (MLP) [18, 19, 21]. We can summarize the attention mechanism in the next three steps.

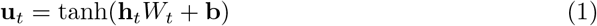

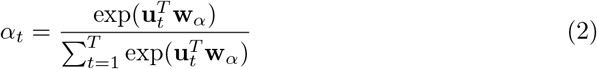

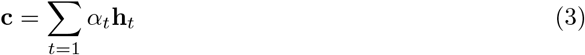

Vectors **h**_*t*_ ∈ ℝ^*H*^ (for *t* ∈ [1, *T*]) are a sequence of hidden representations obtained by a recurrent architecture from an input sequence of events, such as health service claims or visits. These representations are fed to a one-layer MLP with hyperbolic tangent activation to obtain **u**_*t*_ ∈ ℝ^*U*^, a hidden representation of **h**_*t*_ (Eq. 1). Then, a relevance measure of each element in the sequence (*α*_*t*_) is estimated with a Softmax-activated layer (Eq. 2). The weight matrix *W*_*t*_ ∈ ℝ^*H×U*^ and the weight vector **w**_*α*_ ∈ ℝ^*U*^ are jointly learned in the training process. Finally, a context vector **c**can be estimated by computing a weighted sum of the hidden representations **h**_*t*_, with weights *α*_*t*_ (Eq. 3). The context vector can then be further transformed by deeper layers, in order to better approximate the target label [19, 20]. A schematic representation of the attention mechanism is summarized in Fig. 2.

**Fig 2.**
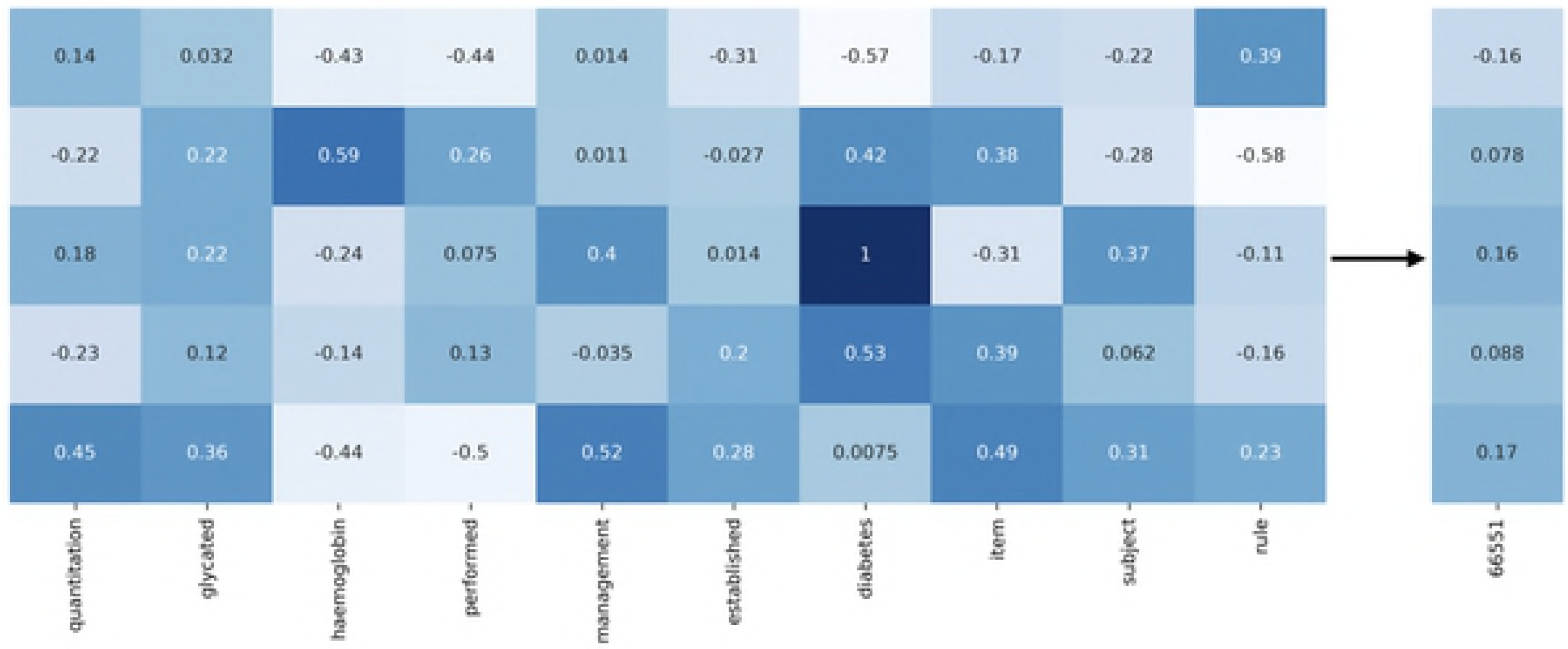
Neural attention model. A visual representation of the attention mechanism for sequence classification. When *λ* = 1 this corresponds to a standard bidirectional attention model for sequence classification, whereas when *λ*≠ 1 the timespan sequence *τ*_1_,…, *τ*_*T*_ can guide the model to focus on the most relevant elements of the sequence. We call `Tangle` the case in which the value of *λ* is jointly learned during the training process. The dashed line highlights the timestamps attention guiding mechanism.

The use of neural attention models for health-related predictions is extensively explored in literature. For instance, in [21] the authors introduce Dipole, a bidirectional recurrent architecture that exploits neural attention to perform sequential EHR forecasting. Moreover, in [16] the authors propose `GRAM`, a graph-based neural attention model that exploits medical ontologies to guide the *α*-estimation step. Finally, in [20] the authors introduce `RETAIN`, a neural attention model for prediction from sequential EHR. `RETAIN` is probably the most relevant work for our purposes. Such model uses two attention levels which separately learn two attention weights vectors that are eventually combined to obtain the context vector. This model achieves good performance when used to predict future diagnosis of heart failure. Although, as the authors claim, it is not capable of exploiting the information hidden in the timestamps of each element of the sequence, which are simply concatenated to each visit embedding ^2^.

## Data

In this work, we analyzed seven years of deidentified records (2008-2014) of the 10% publicly available linked sample of Medicare Benefits Schedule (MBS) and Pharmaceutical Benefits Scheme (PBS) electronic databases of Australia [22]. MBS-PBS 10% sample dataset keeps track of Medicare services subsidised by the Australian government providing information on about 2.1 millions of Australians, who are representative of the full population [23]. The two datasets are linked, meaning that it is possible to track over time the same individual across MBS and PBS claims. MBS-PBS 10% dataset also keeps track of other information such as patients’ gender, state of residence and year of birth. PBS data consist of pharmacy trasactions for all scripts of drugs of the PBS schedule which are dispensed to individuals holding a Medicare card. In PBS, diabetes controlling drugs are identified by 90 item codes grouped in two categories: *insulin and analogues* and *blood glucose lowering drugs, excl. insulins*, the latter including metformins. A difficulty that arises when using this dataset to extract MBS claims trajectories for a given subject is a rule called *episode coning*. According to it, only the items corresponding to the three most expensive pathologies in an episode of care can be contextually claimed and, therefore, can be extracted from the dataset. The rule does not apply to pathology tests requested for hospitalised patients or ordered by specialists.

## Methods

This section provides a detailed definition of the experimental designed followed for the analysis of MBS-PBS 10% dataset, as well as an accurate description of model development, validation and comparison.

### Data preprocessing and representation

In this work, we used PBS data to extract the subject IDs corresponding to the population of interest. We first identified all the subjects that make habitual use of diabetes-controlling pharmaceuticals such as: *Insulins*, *Biguanides*, *Sulfonamides* and so on. Moreover, as PBS did not record medications of non-concessional subjects before 2012, we restricted our analysis to subjects having a concessional card which is used at least for the 75% of the observational years and, in such time interval, for at least 75% of their annual PBS items claims. Such inclusion criteria allowed us to focus on a stable cohort of concessional individuals with diabetes. From this cohort we also identified and excluded records corresponding to subjects with gestational diabetes.

Finally, we labeled with *y*_*i*_ = 1 all the subjects that at first were using only Metformin to manage their diabetes and successively were prescribed to a second-line therapy based on a different drug. This includes both patients that stopped using Metformin at all and patients that associated it with another drug. Conversely, we labeled with *y*_*i*_ = 0 patients that, during the observational time, did not change their Metformin-based diabetes control therapy. This led us to an imbalanced dataset with 26753 subjects which ≈22% are positive.

For each subject in our cohort we used the MBS dataset to extract the corresponding trajectory of Medicare service claims, which can be represented as the following sequence of tuples

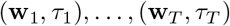

where **w** ∈ ℝ^*V*^ and *τ* ∈ ℕ. The vectors **w**_*t*_ are *V* -dimensional OHE representations of MBS items and the scalars *τ*_*t*_ represent the timespan between two subsequent MBS 159 items, measured in number of days. In our dataset, *V* = 2774 is the vocabulary size (*i.e.* the number of unique MBS items) and *T* = 445 is the sequence length. Sequences shorter than *T* are zero-padded at their beginning, to prevent samples from having inconsistent representations. The first few entries of a prototypical MBS-timespan sequence can look like

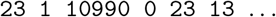

where **w**_1_ = OHE(23), **w**_2_ = OHE(10990), **w**_3_ = OHE(23) while *τ*_1_ = 1, *τ*_2_ = 0 and *τ*_3_ = 13. The 10 most frequent MBS items of our dataset are summarized in Table 1. Dealing with this kind of data, we shall keep in mind that different MBS items may have almost identical meaning. For instance, items 23 and 5020 both apply for general practitioner visits, but the second is dedicated to after-hour attendances. This can be a confounding factor that we will address in the model development process with the help of a pre-trained word embedding.

In order to cope with class imbalance, we matched positive and negatives samples by age (average on the observational time), gender, last pin state and sequence length via Coarsened Exact Matching (CEM) [24]^3^. Table 2 is a summary table of the matched variables statistics before and after CEM matching.

**Table 1.** Summary table of the most frequent MBS items (2.048.502 in total). Items with almost identical meaning are grouped together.

| % | MBS items | Short description |
| --- | --- | --- |
| 0.237 | 10990, 10991 | Management of bulk-billed services |
| 0.187 | 23, 36, 5020, 5040 | General practitioner attendances |
| 0.059 | 73928, 73929, 73938 | Collection of one or more specimens |
| 0.037 | 66503, 66506, 66512, 66515, 66509 | Quantitation of substances in body fluids |
| 0.035 | 74995 | Bulk-billing incentive |
| 0.023 | 65070 | Haematology |
| 0.014 | 10962, 10964 | Podiatric or chiropractic health service |
| 0.014 | 128, 116 | Consultant physician attendances |
| 0.014 | 66551 | Quantitation of HbA1c |
| 0.013 | 105, 108 | Specialist attendances |

**Table 2.** Summary table of the extracted dataset *Pre* and *Post* matching.

|  |  | *Pre* | *Post* |
| --- | --- | --- | --- |
| # Subjects |  | 26753 | 11744 |
| Label (% Class 1) |  | 22.02 | 50.00 |
| AGE (years) |  | 66.15 ± 14.99 | 66.35 ± 11.49 |
| GENDER (% Female) |  | 55.83 | 49.22 |
| SEQUENCE LENGTH (# MBS items) |  | 430.05 ± 364.90 | 347.86 ± 275.31 |
| PIN STATE | % ACT+NSW | 39.49 | 35.87 |
|  | % VIC+TAS | 26.15 | 28.73 |
|  | % WA | 8.67 | 8.65 |
|  | % NT+SA | 8.99 | 9.40 |
|  | % QLD | 16.70 | 17.35 |

### Model development

`Tangle` is a two-inputs/one-output recurrent architecture which, given a set of MBS-timespan sequences, returns the corresponding class probability. A pictorial representation of the model can be seen in Fig. 2. In `Tangle`, the joint MBS-timespan sequence is decoupled in two homogeneous sequences **w**_*t*_ (for *t* = 1, 3, 5,…) and *τ*_*t*_ (for *t* = 2, 4, 6,…) which are used as separate inputs of the network. The vectors **w**_*t*_ are *V* -dimensional OHE representations of MBS items. At the first layer of the network these representations are projected on a *E*-dimensional semantic space, as in Eq. 4, where **x**_*t*_ ∈ ℝ^*E*^ and *W*_*e*_ ∈ ℝ^*V ×E*^.

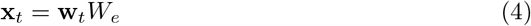

The vocabulary size *V* is defined as the number of unique MBS items observed (plus a dummy entry for the padding value), while the size of the semantic space *E* is a free parameter of the model. In this work we tested two options for the initialization of *W_e_*: uniform random and based on the popular word-embedding GloVe [9]. More details on this second choice will be provided in the next section.

Hidden representations of the two input sequences, **x**_1_,…, **x**_*T*_ and *τ*_1_,…, *τ*_*T*_, are then achieved by two bidirectional LSTM layers [15] (see Eq. 5).

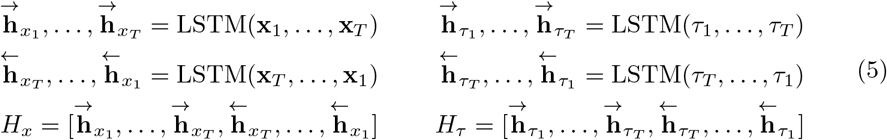

Let *H*_*x*_ ∈ ℝ^*T* ×2*H*^ be the MBS bidirectional hidden representation, where *H* is the number of LSTM units. Similarly, *H*_*τ*_ ∈ ℝ^*T* ×2*H*^ is the bidirectional hidden representation of the timespan sequence. For ease of notation, we define 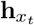 and 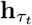 for *t* = 1,…, *T* as generic 2*H*-dimensional vectors belonging to the matrices *H*_*x*_ and *H*_*τ*_, respectively.

The timespan-guided neural attention mechanism adopted in `Tangle` can be described by the following steps.

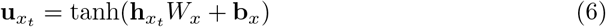

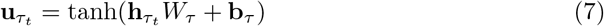

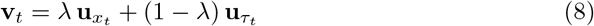

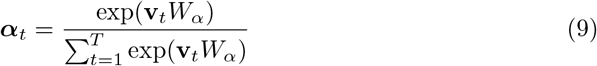

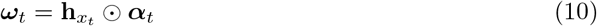

Following the standard attention mechanism, 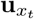 and 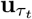 are hidden representations of the sequences 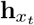 and 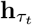 (for *t* = 1,…, *T*). These two vectors are achieved by a one-layer MLP having hyperbolic tangent activation (Eq. 6 and 7). Then, the two hidden representations are merged together in a convex combination **v**_*t*_ ∈ ℝ^*U*^ (Eq. 8), where the mixin parameter *λ* is jointly learned at training time. This is the first novel contribution introduced by the proposed attention mechanism, with respect to the state-of-the-art.

The sequence of **v**_*t*_ is then used to obtain the weights ***α***_*t*_ ∈ ℝ^2*H*^ via Softmax-activated one-layer MLP (Eq. 9). Finally, the attention contribution to each input element ***ω***_*t*_ ∈ ℝ^2*H*^ is expressed as the element-wise product between MBS-sequence hidden representations and the corresponding attention weights (Eq. 10). Interestingly, in our case *W*_*α*_ ∈ ℝ^*U* ×2*H*^, which is the weight matrix of the Softmax layer, plays also the role of projecting the data back to a 2*H*-dimensional space, compatible with LSTM hidden representations. So, each entry of the vectors 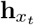 and 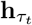 (*i.e.* the output of each LSTM unit) is individually weighted. This is the second original contribution introduced by the proposed attention mechanism with respect to state-of-the-art attention. While the same scalar weight is usually associated to each of the 2*H* entries of the hidden representation **h**_*t*_, `Tangle` is more general as it estimates for each element in the sequence a 2*H*-dimensional attention weights vector.

The context vector c̄ ∈ ℝ^E^ is eventually computed in two steps: first by multiplying along the temporal dimension the contribution matrix

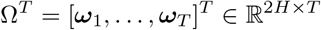

with the input MBS-items sequence matrix

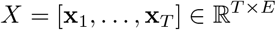

and secondly by average-pooling the 2*H* hidden representations (Eq. 11).

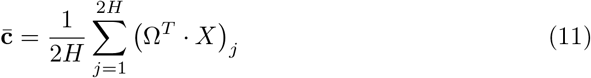

In the proposed architecture, the average context vector **c̄** is fed as input to a two-layers fully connected MLP and trained with Dropout [25]. The first fully connected layer has Rectified Linear Units (ReLu) activation [26], while the output probability is achieved by sigmoid *σ*(·) (Eq. 12).

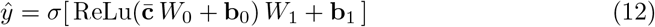

`Tangle` is trained minimizing the Cross-entropy loss (Eq. 13), where *y* ∈ {0, 1} is the binary label associated with the two classes and *N* is the number of samples.

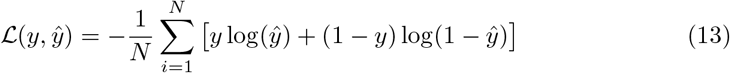

`Tangle` is implemented in Python using Keras [27] and its source code is publicly available on GitHub at https://github.com/samuelefiorini/tangle.

### Embedding weights initialization

As previously anticipated, we need to define a protocol to initialize the embedding matrix *W*_*e*_ (see Eq. 4), which is further optimized in the training phase. The goal of this matrix is to project each MBS item in a semantic space where neighboring points correspond to MBS claims with similar meanings (see Table 1), hence working around the problem of synonym sequence elements.

We first obtained a brief textual descriptions for all the 2774 MBS items by querying the Australian Department of Health website: http://www.mbsonline.gov.au. Then, we cleaned each text corpus from punctuation and stop words and we split the resulting descriptions in 1-grams. For instance, the word list associated to item 66551 is the following.

> [quantitation, glycated, haemoglobin, performed, management, established, diabetes, item, subject, rule]

Then, we associated to each word of the list the corresponding *E*-dimensional `glove.6B` embedding vector, which has 4 × 10^5^ words and it is trained on *Wikipedia 2014 + Gigaword 5* datasets [9]. As of today, `glove.6B` comes in four increasing dimensions: 50, 100, 200, 300. In our experiments we used *E* = 50. Empirical evidences showed that larger embedding dimensions did not significantly increase `Tangle` prediction performance. Finally, we averaged all the single word representations, achieving an *E*-dimensional vector for each MBS item. A pictorial representation of this procedure is depicted in Fig. 3. To demonstrate the effectiveness of our approach, we also tested `Tangle` with uniformly random initialized embedding matrix *W*_*e*_.

**Fig 3.**
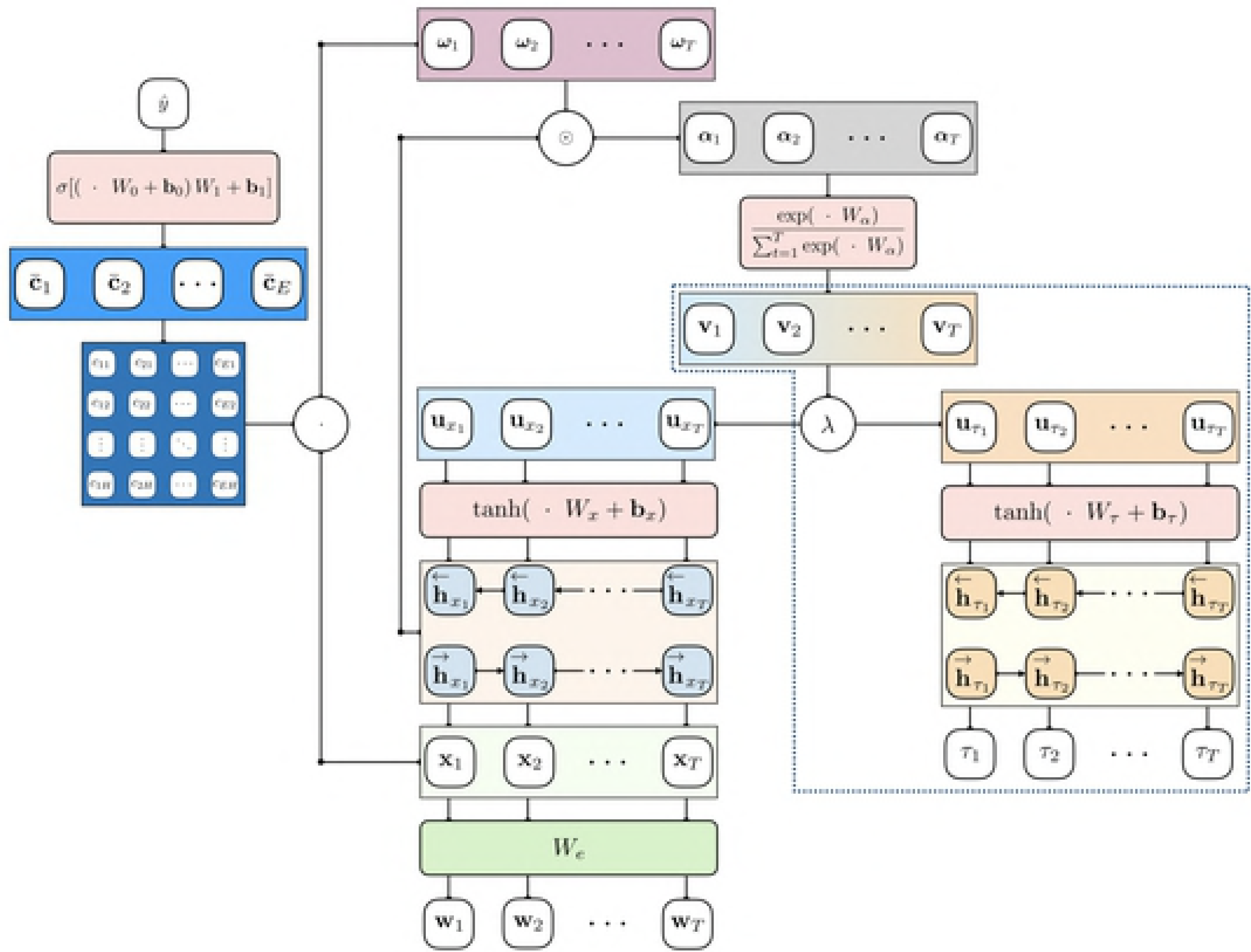
MBS item embedding. A schematic representation of our GloVe-based strategy to achieve meaningful representations of MBS items. To each word of the textual description is associated the corresponding GloVe vector. The final MBS item representation is achieved by averaging.

### Model comparison and analysis

Performance of `Tangle` are evaluated against three different predictive solutions.

1. *ℓ*_1_-penalized LR (see Eq. 14) fitted on a *n*-BOW representation, where *n* controls the number of *n*-grams.

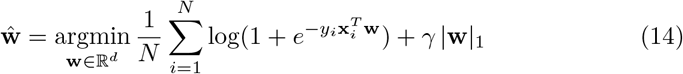 In this case, **x**_*i*_ represents the *n*-BOW representation of the *i*-th patient and *d*, the dimensionality of the LR weights, (**w**) depends on the number of considered *n*-grams.
2. Baseline attentionless recurrent model with bidirectional LSTM (see Fig. 1).
3. State-of-the-art neural attention model with bidirectional LSTM (see Fig. 2).

In order to present a fair model comparison, each tested recurrent model has the same depth, and the only difference is the attention strategy used. Performance of the tested models are evaluated via 10-split Monte Carlo cross-validation [28]. We estimated mean (*µ*) and standard deviation (*σ*) of prediction accuracy, sensitivity, specificity and Area Under the Receiver Operating Characteristics Curve (ROC AUC) [29]. The same 10 Monte Carlo extraction are used for every model. In each Monte Carlo extraction, the matched dataset (with *N* = 11744 samples) is split in two chunks, namely *learning* (60%) and *test* (40%). The learning set is then further split in *training* (90%) and *validation* (10%). This is led us to extract 6341 training, 705 validation and 4698 test samples for each Monte Carlo split. Training sets are used to learn the weights of every model; whereas, validation sets are used by recurrent methods to define the early stopping iteration, and by *ℓ*_1_-LR to optimize the hyperparameter *γ*, which is chosen from a grid of 10 values spanning from 10^*−*5^ to 1 in logarithmic scale. Model predictive performance are then evaluated on each previously unseen test samples.

## Results

We tested three increasing values for *n*: [1, 2, 3]. Choosing *n* = 1 yields the best performance, so results obtained with *n* ≠ 1 are not shown. The grid-search schema used to tune the regularization parameter *γ* of *ℓ*_1_-LR typically resulted in choosing 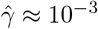. Unpenalized LR was also tested, consistently achieving worse performance. The methods performance is measured in terms of ROC AUC, overall accuracy, sensitivity and specificity [29]. For each performance measure we estimated mean (*µ*) and standard deviation (*σ*) across 10 Monte Carlo samplings. Results of the experiments are summarized in Table 3.

**Table 3.** Summary table comparing the performance of linear and recurrent models. *GloVe initialization of the embedding weight matrix. **Bold** digits highlight best results.

|  | ROC AUC |  | Accuracy |  | Sensitivity |  | Specificity |  |
| --- | --- | --- | --- | --- | --- | --- | --- | --- |
| | $\mu$ | $\sigma$ | $\mu$ | $\sigma$ | $\mu$ | $\sigma$ | $\mu$ | $\sigma$ |
| $\ell_1$-LR 1-BOW | 0.82 | 4.9e-3 | 0.74 | 4.8e-3 | 0.67 | 1.5e-2 | 0.81 | 1.1e-2 |
| Baseline | 0.81 | 8.4e-3 | 0.74 | 7.7e-3 | 0.61 | 4.4e-2 | 0.86 | 4.0e-2 |
| Attention | 0.84 | 1.1e-2 | 0.76 | 1.2e-2 | 0.72 | 4.4e-2 | 0.80 | 5.0e-2 |
| Tangle | 0.87 | 7.8e-3 | 0.78 | 9.9e-3 | 0.71 | 2.6e-2 | 0.85 | 2.7e-2 |
| Baseline* | 0.84 | 9.0e-3 | 0.76 | 9.0e-3 | 0.67 | 5.8e-2 | 0.84 | 5.2e-2 |
| Attention* | 0.86 | 1.2e-2 | 0.77 | 1.1e-2 | 0.71 | 3.9e-2 | 0.83 | 3.9e-2 |
| Tangle* | **0.90** | 6.0e-3 | **0.82** | 8.4e-2 | **0.79** | 3.1e-2 | **0.86** | 3.3e-2 |

Focusing on recurrent methods, `Tangle` outperforms baseline and state-of-the art neural attention architectures. It is interesting to notice how the proposed GloVe-based initialization protocol of the embedding matrix (starred* rows in Table 3) consistently improves on every recurrent model to achieve higher ROC AUC and better classification accuracy. We therefore assume that initializing the embedding weights using GloVe ameliorates the issue of synonym MBS items. Fig. 4 shows the average ROC curve obtained by `Tangle` and *ℓ*_1_-LR that are top and worst performing model, respectively. An intuitive visualization of the discriminative power of the representation achieved by `Tangle` can be seen in the 3D scatter plot of Fig. 5 which was obtained by estimating a 3-dimensional t-SNE embedding [30] on the final sample representation learned by `Tangle`. The figure clearly shows that the learned features are able to discriminate between the two classes, explaining the good performance shown in Table 3.

**Fig 4.**
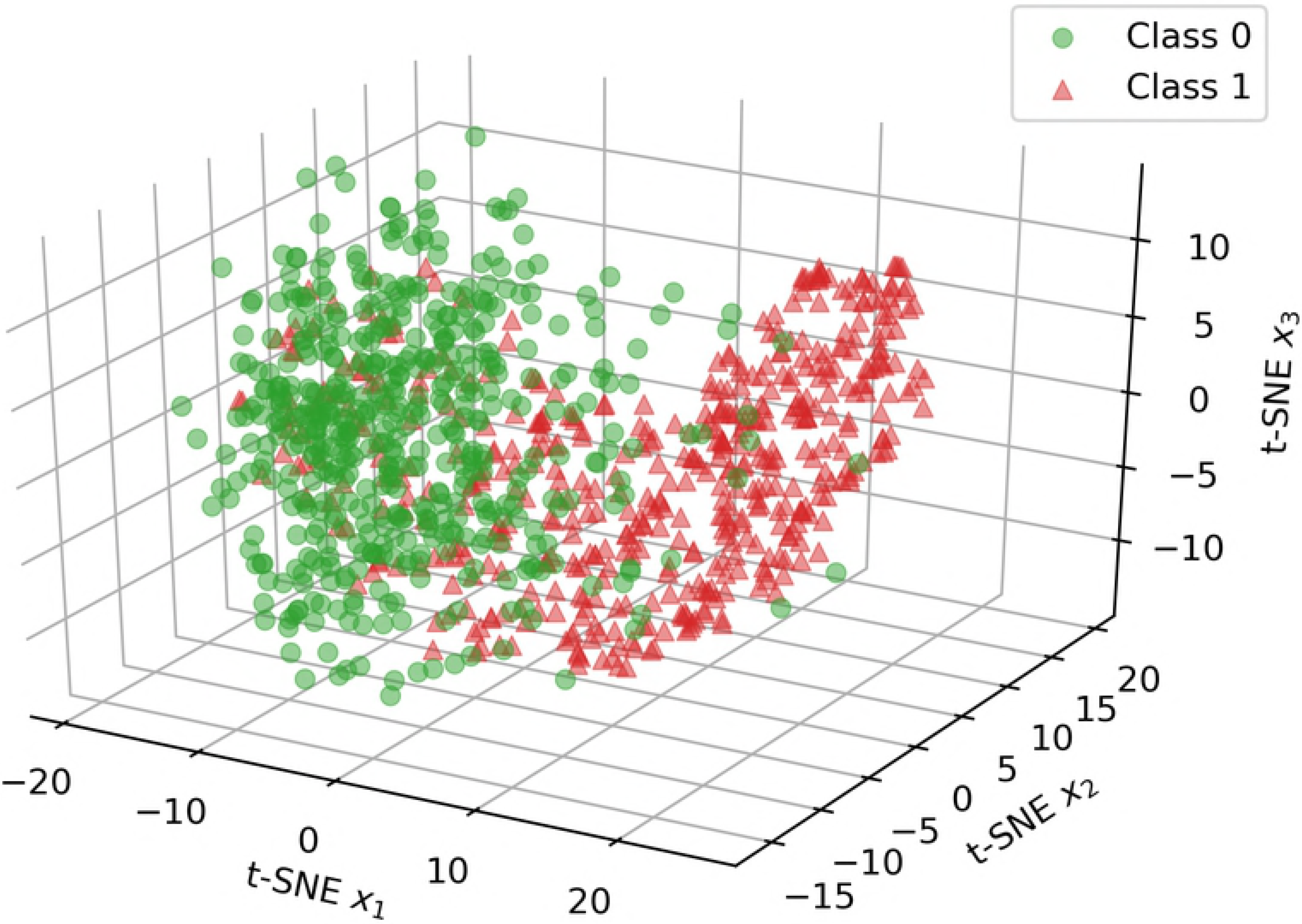
Average ROC curves. ROC curves obtained averaging the 10 Monte Carlo cross-validation iterations for best and worst method: *i.e.*`Tangle` and *ℓ*_1_-LR 1-BOW respectively. Shaded area corresponds to *±*3*σ*, where *σ* is the standard deviation.

**Fig 5.**
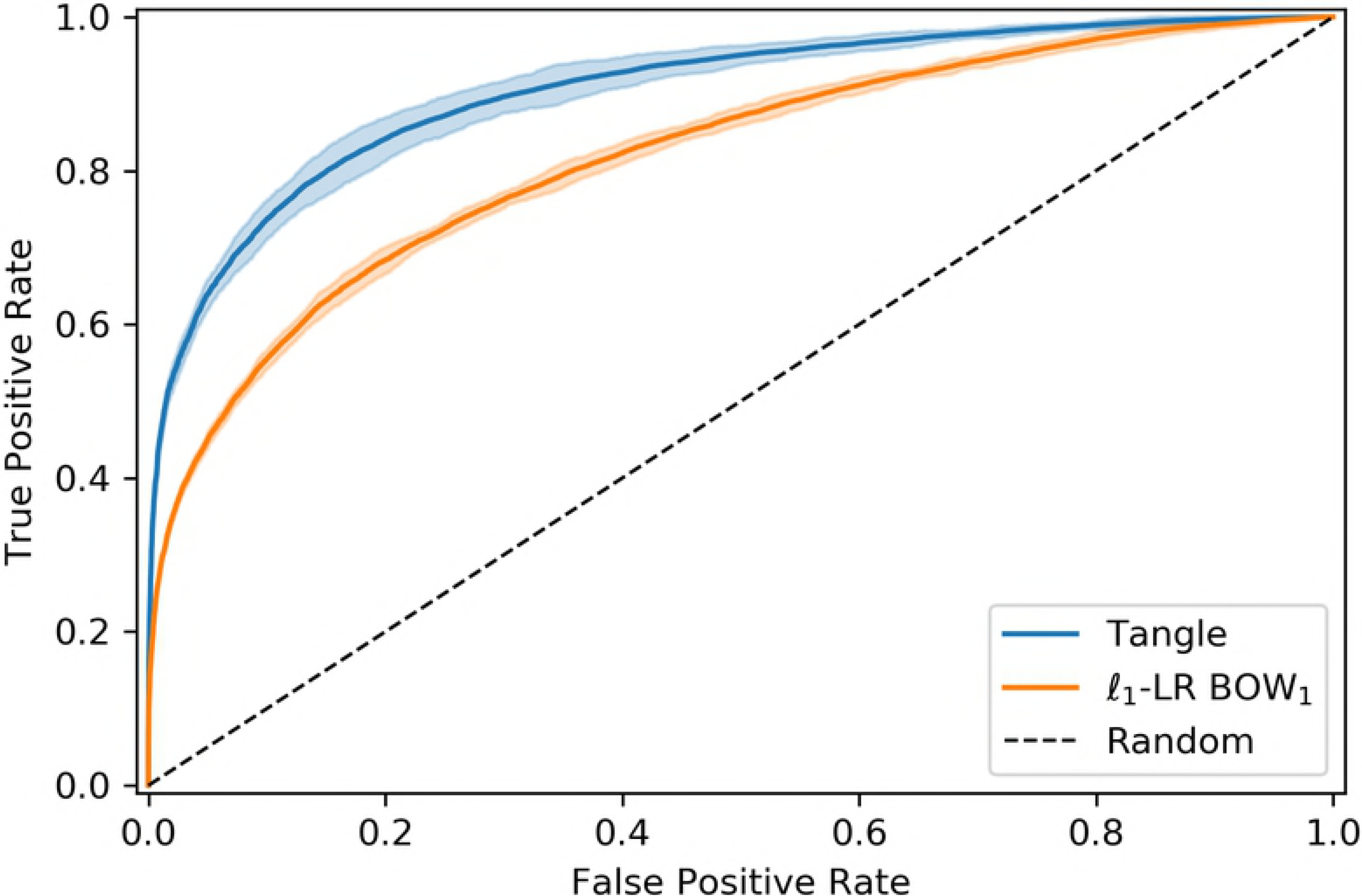
t-SNE embedding. 3D scatter-plot of a random extraction of 500 samples projected on a low-dimensional embedding, estimated by t-SNE [30], from the sample representation learned by `Tangle`. Samples belonging to the two classes, represented with green circles and red triangles, can be seen as slightly overlapping clusters.

A visual representation of the attention contribution estimated by `Tangle` on the test set can be seen in the Manhattan plot of Fig. 6. The horizontal axis corresponds to the MBS items sequence, while their average attention contribution 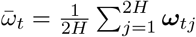 is on the vertical axis. For ease of visualization only the last 250 MBS claims are represented. MBS-items with high attention weight are defined as the ones having 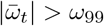 where *ω*_99_ corresponds to the 99-th percentile of the 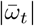 distribution (for *t* = 1,…, *T*). From Fig. 6 we can see that for both classes high attention weights are more frequently falling on the last 13 MBS-items of the sequence, which corresponds to the last 78 days (median value) before the second-line therapy transition. Moreover, we can appreciate how the specific attention weight pattern is different between the two classes.

**Fig 6.**
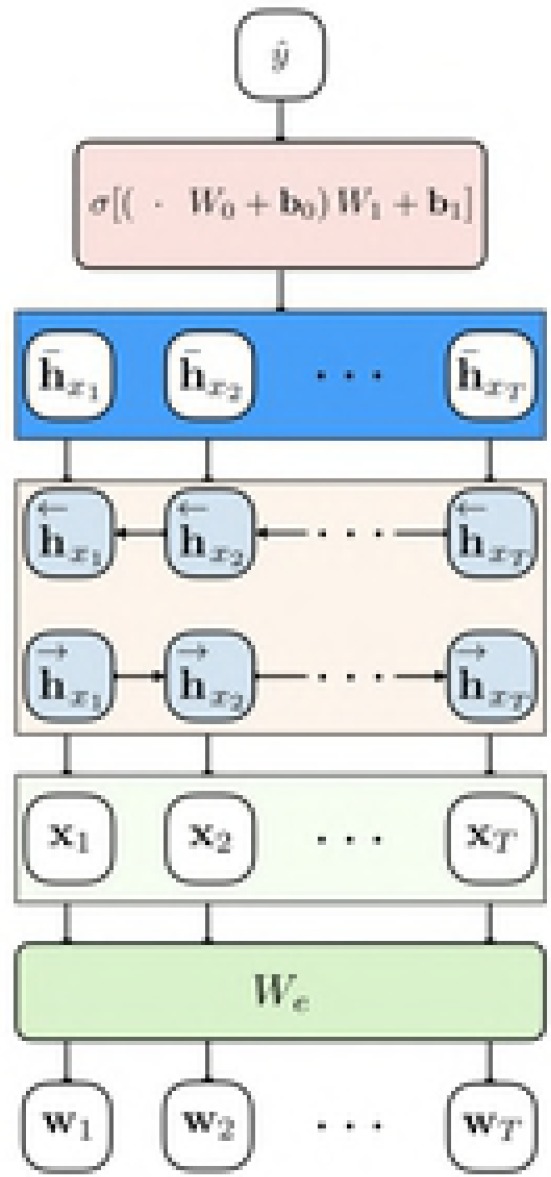
Attention contribution. Manhattan plot of the attention contribution ***ω*** estimated by `Tangle` on the test set. As we can see, the model correctly focuses its attention on the most recent claims, which have nonzero contributions. From this plot we can also appreciate the different representations learned for the two classes.

## Discussion

Our analysis confirms the predictive potential of recurrent models that use neural attention. Interestingly using standard RNNs alone did not substantially outperform simple linear models while requiring a significant computational effort. However, adding the attention mechanism makes the additional computational requirement worth it, since it leads to improved performance. In addition, the proposed timespan-guided attention strategy leads to even better performance, especially if coupled with pre-trained embedding initialization of the weight matrix. Overall, thanks to the available software implementation based on modern deep learning libraries, using `Tangle` does not require significant additional coding effort.

Another advantage of the attention mechanism is that it allows to get an understanding of which portion of the sequence might be more important. For example, in our case we found that the last 13 MBS claims,which take place in ≈78 days, are the most relevant for the current prediction task.

Overall, given that sensitivity and specificity of `Tangle` are at or above 80%, it seems that it could become the basis of an alert system for patient and providers. Clearly, before `Tangle` can be used in practice one would have to understand at which point of the ROC curve of Fig. 4 one should operate. This would require a careful analysis of the relative costs of false positives and false negative alert.

It is important to underscore that there is nothing specific to diabetes in `Tangle`. The modeling strategy and the embedding method could be applied to any problem of sequence classification, providing an easy-to-use method to represent and classify sequences composed of discrete event codes. For example one could apply this method to the analysis of hospital data, where instead of MBS items one has ICD codes, or to more complex data sets, such as the Electronic Health Record collection MIMIC-III [31], that contains clinical codes as well as clinical measures and doctors’ notes.

## Acknowledgments

The authors gratefully acknowledge the support of NVIDIA Corporation with the donation of the Titan Xp GPU used for this research.

## Footnotes

1 Source Australian Government - Department of Health: https://bit.ly/2Njqidp (last visited on January 2019).

2 See RETAIN supplemental material [20]

3 We used the R package cem Version 1.1.19.

## References

1. Australian Government - Australian Institute of Health and Welfare. Diabetes snapshot; 2018. https://www.aihw.gov.au/reports/diabetes/diabetes-compendium/contents/deaths-from-diabetes.

2. Diabetes Australia. Living with diabetes;. https://www.diabetesaustralia.com.au/managing-type-2.

3. Gottlieb A, Yanover, C, Cahan, A, Goldschmidt Y. Estimating the effects of second-line therapy for type 2 diabetes mellitus: retrospective cohort study. BMJ Open Diabetes Research and Care. 2017;5(1):e000435.

4. Kavakiotis, I, Tsave, O, Salifoglou, A, Maglaveras, N, Vlahavas, I, Chouvarda, I. Machine learning and data mining methods in diabetes research. Computational and structural biotechnology journal. 2017;15:104–116.

5. Xing, Z, Pei, J, Keogh, E. A brief survey on sequence classification. ACM Sigkdd Explorations Newsletter. 2010;12(1):40–48.

6. Chollet, F. Deep learning with python. Manning Publications Co.; 2017.

7. Wallach, HM. Topic modeling: beyond bag-of-words. In: Proceedings of the 23rd international conference on Machine learning. ACM; 2006. p. 977–984.

8. Mikolov, T, Sutskever, I, Chen, K, Corrado, GS, Dean J. Distributed representations of words and phrases and their compositionality. In: Advances in neural information processing systems; 2013. p. 3111–3119.

9. Pennington, J, Socher, R, Manning, CD. GloVe: Global Vectors for Word Representation. In: Empirical Methods in Natural Language Processing (EMNLP); 2014. p. 1532–1543. Available from: http://www.aclweb.org/anthology/D14-1162.

10. Friedman, J, Hastie, T, Tibshirani, R. The elements of statistical learning. vol. 1. Springer series in statistics New York; 2001.

11. Breiman, L. Random forests. Machine learning. 2001;45(1):5–32.

12. Freund, Y, Schapire, RE. A decision-theoretic generalization of on-line learning and an application to boosting. Journal of computer and system sciences. 1997;55(1):119–139.

13. Guyon, I, Elisseeff, A. An introduction to variable and feature selection. Journal of machine learning research. 2003;3(Mar):1157–1182.

14. LeCun, Y, Bengio, Y, Hinton, G. Deep learning. nature. 2015;521(7553):436.

15. Hochreiter, S, Schmidhuber, J. Long short-term memory. Neural computation. 1997;9(8):1735–1780.

16. Choi, E, Bahadori, MT, Song, L, Stewart, WF, Sun, J. GRAM: Graph-based attention model for healthcare representation learning. In: Proceedings of the 23rd ACM SIGKDD International Conference on Knowledge Discovery and Data Mining. ACM; 2017. p. 787–795.

17. Cho, K, Van Merriënboer, B, Gulcehre, C, Bahdanau, D, Bougares, F, Schwenk, H, et al. Learning phrase representations using RNN encoder-decoder for statistical machine translation. arXiv preprint arXiv:14061078. 2014;.

18. Bahdanau, D, Cho, K, Bengio, Y. Neural machine translation by jointly learning to align and translate. arXiv preprint arXiv:14090473. 2014;.

19. Yang, Z, Yang, D, Dyer, C, He, X, Smola, A, Hovy, E. Hierarchical attention networks for document classification. In: Proceedings of the 2016 Conference of the North American Chapter of the Association for Computational Linguistics: Human Language Technologies; 2016. p. 1480–1489.

20. Choi, E, Bahadori, MT, Sun, J, Kulas, J, Schuetz, A, Stewart, W. Retain: An interpretable predictive model for healthcare using reverse time attention mechanism. In: Advances in Neural Information Processing Systems; 2016. p. 3504–3512.

21. Ma, F, Chitta, R, Zhou, J, You, Q, Sun, T, Gao, J. Dipole: Diagnosis prediction in healthcare via attention-based bidirectional recurrent neural networks. In: Proceedings of the 23rd ACM SIGKDD International Conference on Knowledge Discovery and Data Mining. ACM; 2017.p. 1903–1911.

22. Australian Government - Department of Health. Public Release of Linkable 10sample of Medicare Benefits Scheme (Medicare) and Pharmaceutical Benefits Scheme (PBS) Data; 2016. http://www.pbs.gov.au/info/news/2016/08/public-release-of-linkable-10-percent-mbs-and-pbs-data.

23. Hajati, F, Atlantis, E, Bell, KJ, Girosi, F. Patterns and trends of potentially inappropriate high-density lipoprotein cholesterol testing in Australian adults at high risk of cardiovascular disease from 2008 to 2014: analysis of linked individual patient data from the Australian Medicare Benefits Schedule and Pharmaceutical Benefits Scheme. BMJ open. 2018;8(3):e019041.

24. Iacus, SM, King, G, Porro, G. Causal inference without balance checking: Coarsened exact matching. Political analysis. 2012;20(1):1–24.

25. Hinton, GE, Srivastava, N, Krizhevsky, A, Sutskever, I, Salakhutdinov, RR. Improving neural networks by preventing co-adaptation of feature detectors. arXiv preprint arXiv:12070580. 2012;.

26. Krizhevsky, A, Sutskever, I, Hinton, GE. Imagenet classification with deep convolutional neural networks. In: Advances in neural information processing systems; 2012. p. 1097–1105.

27. Chollet, F, et al.. Keras; 2015. https://keras.io.

28. Molinaro, AM, Simon, R, Pfeiffer, RM. Prediction error estimation:a comparison of resampling methods. Bioinformatics. 2005;21(15):3301–3307.

29. Everitt, B, Skrondal, A. The Cambridge dictionary of statistics. vol. 106.Cambridge University Press Cambridge; 2002.

30. Maaten, Lvd, Hinton, G. Visualizing data using t-SNE. Journal of machine learning research. 2008;9(Nov):2579–2605.

31. Johnson, AE, Pollard, TJ, Shen, L, Li-wei, HL, Feng, M, Ghassemi, M, et al. MIMIC-III, a freely accessible critical care database. Scientific data. 2016;3:160035.

